# Isotropic expansion of external environment induces tissue elongation and collective cell alignment

**DOI:** 10.1101/619189

**Authors:** H. Koyama, T. Fujimori

**Author notes:** Correspondence: H. Koyama.

## Abstract

Cell movement is crucial for morphogenesis in multicellular organisms. Growing embryos or tissues often expand isotropically, i.e., uniformly, in all dimensions. On the surfaces of these expanding environments, which we call “fields,” cells are subjected to frictional forces and move passively in response. However, the potential roles of isotropically expanding fields in morphogenetic events have not been investigated well. In this study, we mathematically analyzed the effect of isotropically expanding fields using a vertex model, a standard type of multi-cellular model. We found that cells located on fields were elongated along a similar direction each other. Simultaneously, the cell clusters were also elongated, even though field expansion was absolutely isotropic. We then investigated the mechanism underlying these counterintuitive phenomena. In particular, we asked whether elongation was caused by the properties of the field, the cell cluster, or both. Theoretical analyses involving simplification of the model revealed that cell clusters have an intrinsic ability to asymmetrically deform, leading to their elongation. Importantly, this ability is effective only under the non-equilibrium conditions provided by field expansion. This may explain the elongation of the notochord, located on the surface of the growing mouse embryo. We established that passive cell movement induced by isotropically expanding external environments can contribute to both cell and tissue elongation, as well as collective cell alignment, providing key insight into morphogenesis involving multiple adjacent tissues.

**Statement of Significance:** It is a central question of developmental biology how the symmetric shapes of eggs can develop the asymmetric structures of embryos. Embryos expand through their growth. Simultaneously, elongation of tissues such as the notochord occurs, which is fundamental phenomena of morphogenesis. However, possible relationships between tissue elongation and the expansion of embryos have not been investigated well. Here we mathematically present that, even if the expansion is isotropic, tissues located on the embryos are asymmetrically deformed by the expansion, resulting in elongation. We generalize the effect of expanding environments on tissue elongation through model reduction and uncover the mechanism underlying elongation. This process can be a novel key piece for symmetry breaking of embryos, together with previously established morphogenetic processes.

## Introduction

In multicellular organisms, tissues and organs assume many different shapes, and cells exhibit many distinct configurations. These features are generated through developmental processes including cell migration, growth, differentiation, and tissue deformation. Cells have intrinsic features that cause them to migrate in developing tissues, and often undergo directional migration to form a certain tissue morphology or cell configuration. The mechanisms and outcomes of these cell-autonomous migrations have been studied extensively. For instance, cell-autonomous migration in a specific direction causes remodeling of cell-cell junctions, accompanied by exchanges of cellular neighbors, and subsequent convergent extension, resulting in elongation of the tissue (1–4). Thus, these cells have intrinsic abilities to elongate the tissues. By contrast, cell movement can also be passively affected by extrinsic mechanical forces (5). Tensile or frictional forces can be exerted by tissues in contact with one another (6–8). These extrinsic forces are involved in tissue deformations, cell shape changes, collective cell alignment, and patterns of epithelial cell packing (5–9). In these cases, the cells have no intrinsic abilities to deform the tissues.

Cell elongation is accompanied by cell movements and tissue deformations. In several tissues, many cells collectively elongate in the same direction, leading to generation of aligned cell arrays that are involved in morphogenesis and the formation of planar cell polarity (PCP) in epithelia (6, 10, 11). Generation of these arrays can be achieved by active cell deformation or extrinsic mechanical forces. During tissue elongation involving convergent extension, cells actively elongate toward the center line of the elongating tissue; consequently, the direction of cell elongation becomes perpendicular to that of tissue elongation (1, 2, 11, 12). By contrast, during passive elongation of tissues or cells induced by extrinsic mechanical forces, the direction of elongation is usually correlated with that of the extrinsic forces (6, 7).

Although elongation of tissues and cells has been extensively investigated, we discovered a novel type of tissue elongation induced by extrinsic mechanical forces, in which the correlation between directions of tissue/cell elongation and extrinsic forces is lost (13). A mouse post-implantation embryo grows and expands like an expanding rubber balloon, driven by an increase in the volume of the inner cavity. At the same time, the notochord, which is located on the surface of the embryo, elongates (Fig. 1A and B). We showed that in contrast to notochord elongation in other species, which is induced by active cell movement (1, 2, 11), elongation of the mouse notochord is driven, at least in part, by the passive mechanical effects of embryonic expansion (13). Because the expansion of the embryo is almost isotropic, like a spherically expanding balloon, the direction of the extrinsic forces provided from the expanding embryo are expected to be unbiased and uncorrelated with the direction of notochord elongation (Fig. 1B)(13). Furthermore, the cells are also elongated in a direction parallel to that of notochord elongation (Fig. 1A) (13), in contrast to the situation in other elongating tissues, including the notochords of other species, in which the direction of cell elongation is perpendicular to that of tissue elongation (1, 11, 12). Thus, the outcomes of tissue and cell elongation on an isotropically expanding embryo are counterintuitive and difficult to predict, and it remains poorly understood how tissues and cells can deform asymmetrically, resulting in elongation. In general, fields undergoing isotropic expansion in two or three dimensions arise in growing embryos and tissues, or in local regions of tissues, including epiboly during fish embryogenesis, development of the limb bud, etc. (Fig. 1B and E) (1, 14, 15). These observations inspired us to theoretically generalize the effect of isotropically expanding fields on tissue and cell elongation and investigate the underlying mechanisms. In this report, we provide a theoretical explanation of tissue elongation caused by passive/non-cell-autonomous movement on isotropically expanding fields, and describe the influence of these fields on collective cell elongation and alignment.

**Figure 1:**
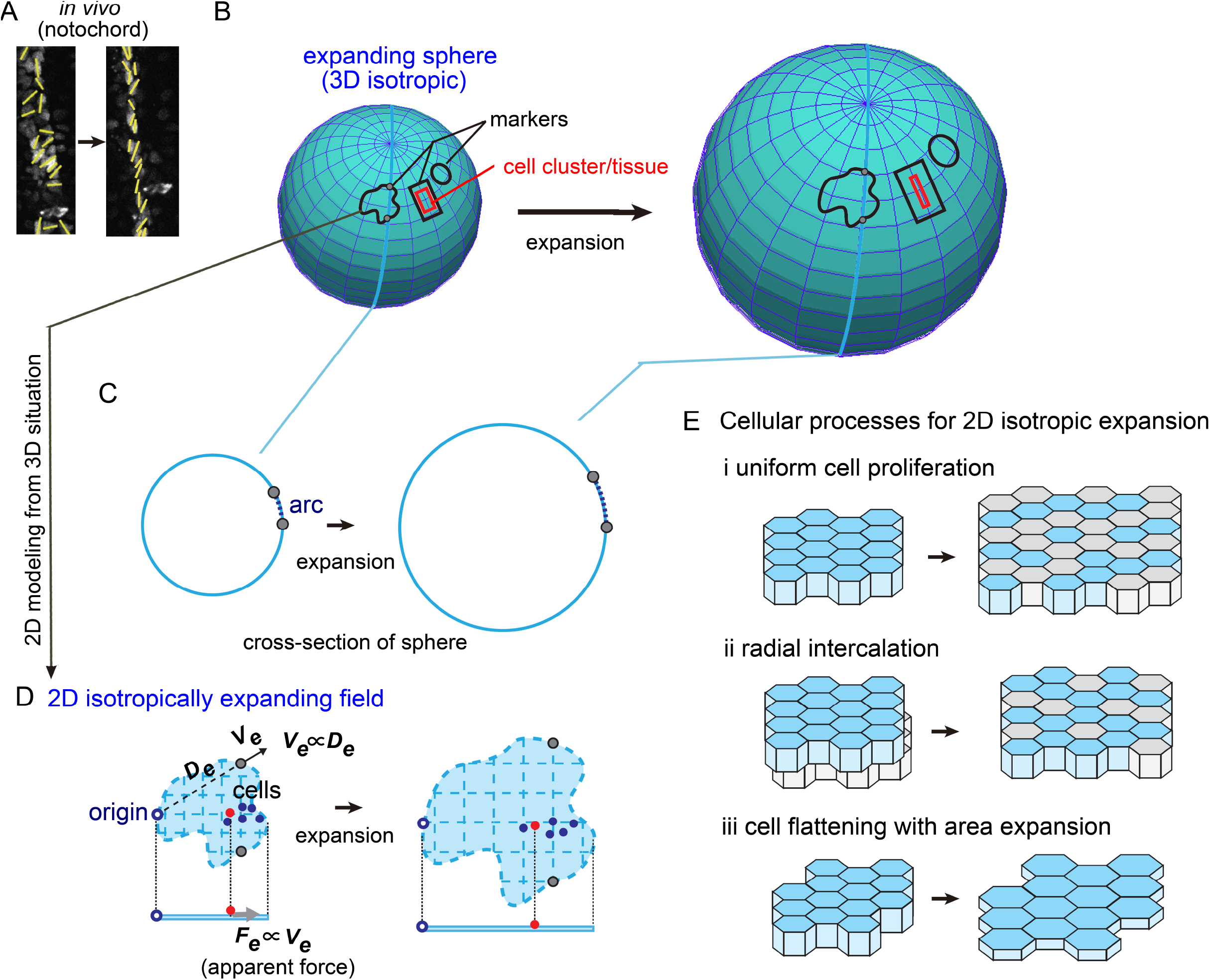
Cell cluster located on an isotropically expanding field. A) Notochord elongation in mouse embryo. Yellow lines indicate orientation of each cell. The width of the notochord decreases, so the tissue must elongate. Images were adapted from Imuta et al (13). B) Schematic illustration of a cell cluster or tissue on an expanding sphere. An expanding sphere is three-dimensionally isotropic, so any regions on the surface expand while maintaining their original shape, other than their size (three black markers). By contrast, a cell cluster on the surface does not maintain its original shape and deforms (red rectangle). C) Cross-section of the sphere in B. Two points on the surface are shown in B and C (gray circles). The distance between two arbitrary points along the surface increases at a rate proportional to the arc length. D) Two-dimensional modeling of an isotropically expanding field. The three-dimensional situation shown in B and C is approximated in two dimensions. An isotropically expanding field (light blue plane) and cells/objects (filled circles) located on it are shown. If an origin (open blue circle) is defined, a point (gray circle) on the field moves away from the origin at a velocity (*V_e_*) proportional to its distance (*D_e_*) from the origin. A cell (red filled circle) experiences an apparent force (*F_e_*) from the field, which is proportional to *V_e_* at the cell’s position. Top and side views are shown with the direction of the apparent force exerted on the red object. E) Examples of cellular processes provoking isotropic expansion of fields in two-dimensional situations. Cells are presented as columnar polygons. Examples of uniform cell proliferation, radial interaction, and cell flattening with area expansion are shown (i–iii). The overall shape of the cell sheet, other than its size, is conserved. i) Newly generated cells are shown in gray. ii) Two or more cell layers become a single cell layer. The gray cells are radially intercalated into the layer of blue cells. iii) Cells are flattened, and their upper surface area increases.

## Methods

### Definition of patterns

Asymmetry index (*AI*) was defined as follows; *AI* = *L_axis_*/*D_circle_*, where *L_axis_* is the length of the longest axis of a cell cluster, and *D_circle_* is the diameter of a circle with the same area as the cell cluster. Thus, *AI* of a circle is 1.0.

### Calculation of morphologies and directions of cells and cell clusters

Areas of cell clusters were measured using the ImageJ software. The lengths of the long and short axes of cell clusters and individual cells were measured as Feret and MinFeret in ImageJ, respectively. The elongation direction of cell clusters and cells were also measured by using FeretAngle in “Fit ellipse” of ImageJ.

## Results

### Mathematical model of an isotropically expanding field

A mouse post-implantation embryo is ovoid or cylindrical during post-implantation (embryonic day 7.5–8.5), and grows while maintaining its overall shape (13). The embryo contains a few cavities filled with liquid, which expand during the growth of the embryo. Thus, the embryo and its cavities resemble, respectively, the surface and inner air of an expanding rubber balloon. The notochord is located on the surface of the embryo, and elongates as the embryo grows (Fig. 1A and B, red cell cluster/tissue).

To simplify our conceptualization of the growing embryo, we approximated its shape as a sphere (Fig. 1B). On the surface of an expanding sphere, the distance between two arbitrary points increases at a rate proportional to their distance (Fig. 1B; e.g., the distance between two gray circles). Note that this rule also holds true if distance is calculated as the length along the curved surface of the sphere (Fig. 1C, arc length between two gray circles). Consequently, shapes on the surface are preserved (except for their sizes) during expansion (Fig. 1B, black markers). In addition, this rule is true on surfaces of an expanding ovoid object, and indeed objects of any shape. This prompted us to further simplify the situation: instead of an expanding curved three-dimensional surface, we assumed an isotropically expanding two-dimensional plane on which the distance between two arbitrary points increases at a rate proportional to their distance (Fig. 1D; e.g., the distance between two gray circles), as described in our earlier study (13). This situation can also arise in a two-dimensional tissue, like a cell sheet, in which spatially uniform cell proliferation, radial intercalation (16, 17), or cell flattening occurs (Fig. 1E)(1), implying that our assumption has generality.

To mathematically model an isotropically expanding two-dimensional plane (or field), it is convenient to define an origin in a two-dimensional x-y plane (Fig. 1D, origin). If one assumes that a point on the field moves away from the origin with a velocity (*V_e_*) proportional to the distance (*D_e_*) between the point and the origin (Fig. 1D, *V_e_* ∝ *D_e_*), one can show that the distance between any two arbitrary points (Fig. 1D; e.g., two gray circles) on the field increases at a rate proportional to the distance. Therefore, under this assumption, an isotropically expanding field can be correctly modeled, as confirmed by simulation results described below (Fig. S1).

### Mathematical model of cells on an isotropically expanding field

In our previous report, to simulate deformation of a cell cluster, we constructed a grid-based model (13), which will be discussed below (Fig. 3A). However, our previous model did not correctly evaluate deformation of individual cells because it did not consider cell neighbor exchange (Fig. 2A, cell junctional remodeling). To simulate deformation of both cell clusters and individual cells, we employed a vertex model, a standard and more realistic multicellular model (Fig. 2A) (18). In this model, each cell is modeled as a polygon in which the line tensions between cell–cell boundaries and area conservation of each cell are considered as follows:

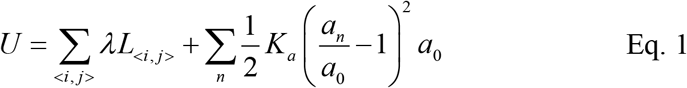

where *U* is the mechanical potential energy of the whole system, and *λ* and *L_<i,j>_* are the line tension and length of cell–cell boundaries between adjacent vertices *i* and *j* (Fig. 2A), respectively (19). The line tensions are derived from contractile forces of cell-cell boundaries, etc. *a_n_* is the area of cell *n. a*_0_ and *K_a_* are a preferred area of the cell and the coefficient of area elasticity, respectively. The forces exerted on each vertex are calculated as follows:

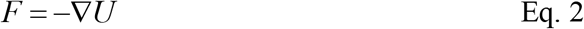

where ∇ is the nabla vector differential operator at each vertex. Importantly, we also considered a type of cell neighbor exchange called a T1 transition, which usually occurs during tissue elongation accompanying convergent extension, and is essential for evaluating cell deformation (Fig. 2A, cell junctional remodeling) (18).

**Figure 2:**
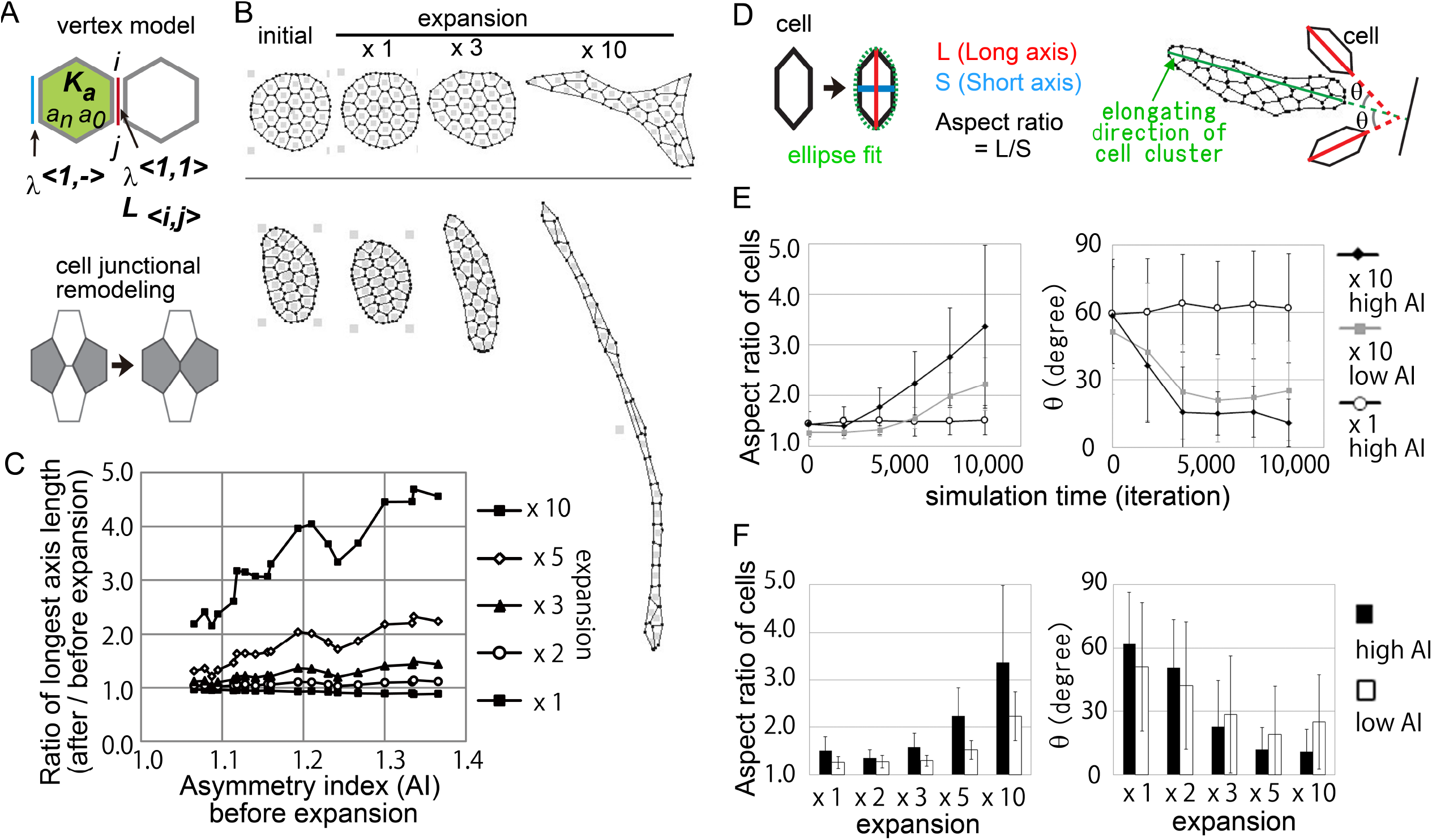
Cell cluster and cell elongation in vertex model. A) Schematic representation of the vertex model and cell junctional remodeling. Line tension *λ* between cells (*λ*^<1,1>^) and on outer boundaries (*λ*^<1,−>^), along with area elasticity (*K_a_*), are shown. *L_<i,j>_* is the length between vertices *i* and *j*. *a_n_* is the area of *n*th cell, and *a_0_* is the preferred or natural area of the cell. Cell junctional remodeling induces cell neighbor exchange. B) Examples of simulations in the vertex model. Two initial cell configurations, one nearly symmetric and another slightly asymmetric, are shown in the first column (initial). During simulations, the fields were expanded by 1-, 3-, or 10-fold as described. The conditions of the simulations were as follows: *λ*^<1,1>^ = 0.4, *λ*^<1,−>^ = 1.0, *K_a_* = 6.0, *a_0_* = 1.0, *μ* = 100.0, simulation time = 2,000 [A.U.]. Simulation programs were written in the C language. C) Initial configuration-dependent rate of elongation. Various initial configurations with different Asymmetry Indices (*AI*) (defined in Methods) were used, and changes in the length of the longest axis were analyzed after the indicated degrees of expansion (× 1, 2, 3, 5, and 10). D) Schematic representation of cell elongation and its orientation. In the left panel, elongation of each cell (polygon) was evaluated by the aspect ratio calculated when the cell is fitted to an ellipse (green broken line). The long and short axes are indicated by red and blue lines, respectively. In the right panel, the orientation of each cell relative to the tissue elongation direction was calculated. The tissue elongation direction is indicated by a green line, and the longer axes of two polygonal cells (enlarged views) are shown as red lines. The angles (θ = 0–90°) between the green and red lines were defined as the orientations of individual cells. E) Aspect ratios of cells and θ in cell clusters during simulations. The simulation data were adopted from B as described in the main text. Each cell cluster contains 26 cells. Error bars are standard deviations. When cells are randomly elongated, the average value of θ is expected to be 45°. F) Aspect ratios of cells and θ in cell clusters under various expansion rates. In addition to the simulation data from B, data obtained with expansion rates of ×2 and ×5 are also shown.

The cells modeled above were placed on a field. We assumed that frictional forces could be exerted between the vertices of the cells and the field, and that the motions of the vertices were damped by this friction. The damping was assumed to be large enough to ignore the viscosity of the medium and inertial effects. In general, when the motions of vertices are damped by friction or viscosity of surrounding tissues, the following relationship is typically assumed (2, 4, 18, 20): *F*=*μV*, where *V* is the velocity of a vertex, and *μ* is the coefficient of the friction or viscosity. Thus, in the absence of field expansion, the motion of the vertex can be written as *V*=*F*/*μ* if neither thermal fluctuations nor cellular noise are considered, as in the simplest vertex model (18, 20). In the presence of field expansion, we assumed that the velocity is modified as *V*=*F*/*μ* + *V_e_*, where *V_e_* is the velocity of the expanding field at the vertex, as previously defined in Figure 1D. Therefore, we can write

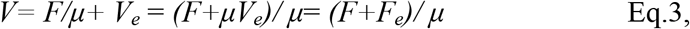

where *F_e_* corresponds to an “apparent” force exerted by the field when a point on the field is defined as the x-y origin (Fig. 1D). Because *V_e_* ∝ *D_e_*, we can write

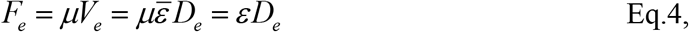

where 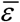 and *ε* are coefficients with different dimensions. Therefore, the apparent force increases when the vertices are located far from the x-y origin, causing the vertices to move rapidly away from the origin. This model was simulated using the Euler method.

Next, we sought to confirm that the isotropic features of the expanding field are correctly expressed by our model. First, we set the cell-intrinsic forces such as line tensions as absent (*F* = 0), and then performed simulations. The movements of the vertices absolutely coincided with the expansion of the field (*V* = *V_e_*), causing no deformation except for area expansion of the cell cluster (Fig. S1A). Next, we set the cell clusters at various positions on the expanding field, e.g., near or far from the x-y origin. The deformation patterns of the cell clusters were absolutely identical among the cell clusters (Fig. S1B). This was also the case when the cell-intrinsic forces were set as present (*F* ≠ 0) (Fig. S1C). In addition, when the x-y origin was set at different positions, the deformation patterns of the cell clusters were not affected (Fig. S1D). These results indicated that, although the apparent forces exerted by the field were locally different, our model achieved an absolutely isotropic field.

### Elongation of cell clusters

Using our vertex model, we investigated whether a cell cluster or individual cell would be elongated during isotropic expansion of the field. Under conditions of no field expansion, cell clusters tended to form a circular configuration similar to typical endothelial cell clusters, decreasing their initial asymmetry (Fig. 2B, bottom row; expansion ×1). Thus, the cell clusters did not elongate autonomously. In other words, the cell clusters did not seem to have intrinsic features that cause them to elongate. By contrast, under conditions of an expanding field, a slightly elongated cluster of cells became more elongated (Fig. 2B, bottom row; expansion ×3 and ×10), consistent with our previous conclusions (13). Thus, the asymmetry of cell cluster shape was enhanced by field expansion. Interestingly, cell neighbor exchanges were extremely frequent, leading to an elongated column of nearly single-cell width (expansion ×10), consistent with the situation in mouse notochord *in vivo* (Fig. 1A) (13). A symmetric, almost circular initial configuration yielded a distorted shape, i.e., the initial symmetry was broken (Fig. 2B, upper row; expansion ×10). To quantify the increase in asymmetry of cell clusters, we compared the length of the longest axis of a cell cluster before and after the field expansion for initial configurations with various asymmetry index (*AI*) values (*AI* = 1.0 if the cell cluster is circular, as defined in Materials & Methods). The length of the longest axis was increased to a greater extent for initial configurations with high *AI* values than for those with low *AI* values (Fig. 2C), explicitly demonstrating the dependence of elongation/asymmetry enhancement on the initial configuration.

### Elongation and alignment of cells

Next, we evaluated deformations of individual cells. In the elongated cell clusters, individual cells were also elongated in almost the same direction (Fig. 2B, bottom row; expansion ×10). Elongation was quantified as the aspect ratio of each cell (Fig. 2D, aspect ratio). We then analyzed the cell cluster shown in the bottom row of Figure 2B, for which the initial cell configuration exhibited a high *AI* value. The aspect ratios of the cells increased temporally in an elongating cell cluster (Fig. 2E, left panel, “×10 high *AI*” which corresponds to Figure 2B bottom row, expansion ×10), but not in a non-elongating cell cluster (Fig. 2E, left panel, “×1 high *AI*,” which corresponds to Figure 2B bottom row, expansion ×1). Even in a cell cluster exhibiting slight elongation, the aspect ratios were elevated (Fig. 2E, left panel, “×10 low *AI*” which corresponds to Figure 2B upper row, expansion ×10). Analyses of conditions with various expansion rates revealed that the increase in aspect ratios depended on the expansion rates (Fig. 2F, left panel; ×1, 2, 3, 5 vs. 10). These results indicate that isotropically expanding fields can induce cell elongation.

We next investigated the effect of field expansion on the collective behaviors of cells. To this end, we analyzed the orientations of cell elongation relative to the elongation direction of the cell cluster (Fig. 2D, θ, which is defined from 0 to 90 degrees). The angle θ relative to the long axis of the cell cluster decreased temporally in an elongating cell cluster (Fig. 2E, right panel, “×10 high *AI*” and “×10 low *AI*”) but not in a non-elongating cluster (Fig. 2E, right panel, “×1 high *AI*”). Thus, the orientations of cell elongation were parallel to the direction of elongation of the cell cluster, consistent with the situation in the mouse notochord (Fig. 2E, right panel vs. 1A and Figure 2F in (13)). Moreover, the standard deviation of θ was reduced in elongating cell clusters, suggesting that cell elongation orientations became aligned with each other (Fig. 2E, right panel, “×10 high *AI*” and “×10 low *AI*”). Analyses of conditions with various expansion rates revealed that the decreases in θ and alignments of cell elongation orientations depended on the expansion rate (Fig. 2F, right panel; ×1, 2, 3, 5, and 10). These findings indicate that isotropically expanding fields can contribute to collective alignment of cell elongation orientations toward the elongation directions of cell clusters.

### Simplification of the model

We showed that cell clusters are elongated on isotropically expanding fields, although the cell clusters do not seem to have intrinsic features that cause them to elongate and the fields do not have anisotropic features (Fig. S1). What mechanisms underlie this counterintuitive phenomenon? Where are the anisotropic or asymmetric features hidden in this system? More specifically, are elongations of cell clusters and individual cells primarily caused by the properties of the field or the cell cluster, or by their combined effects?

To address these questions, we sought to analytically dissect the roles of the properties of the field and cell clusters on the elongation of the clusters. For this purpose, the vertex model was too complicated, so we tried to simplify it. First, we modeled a cell cluster as a grid, as described previously (Fig. 3A) (13), in which cell neighbor exchanges were not permitted. The results obtained from simulations using grid model are fundamentally similar to those obtained using the vertex model: in both models, cell clusters were elongated on isotropically expanding fields. The grid model was composed of *i* × *j* cells, as shown in Figure 3A. Apparent forces from the expanding field were considered to be exerted at each vertex in a manner similar to the vertex model (Fig. 3A, middle panel). Line tensions between cell–cell boundaries and area conservation of each cell were also implemented similarly to the vertex model (Fig. 3A, right panel; *λ* and *K_a_* are the line tensions and the coefficient of area elasticity, respectively). In Figure 3A, the x- and y-axes were set along the edges of the cluster.

**Figure 3:**
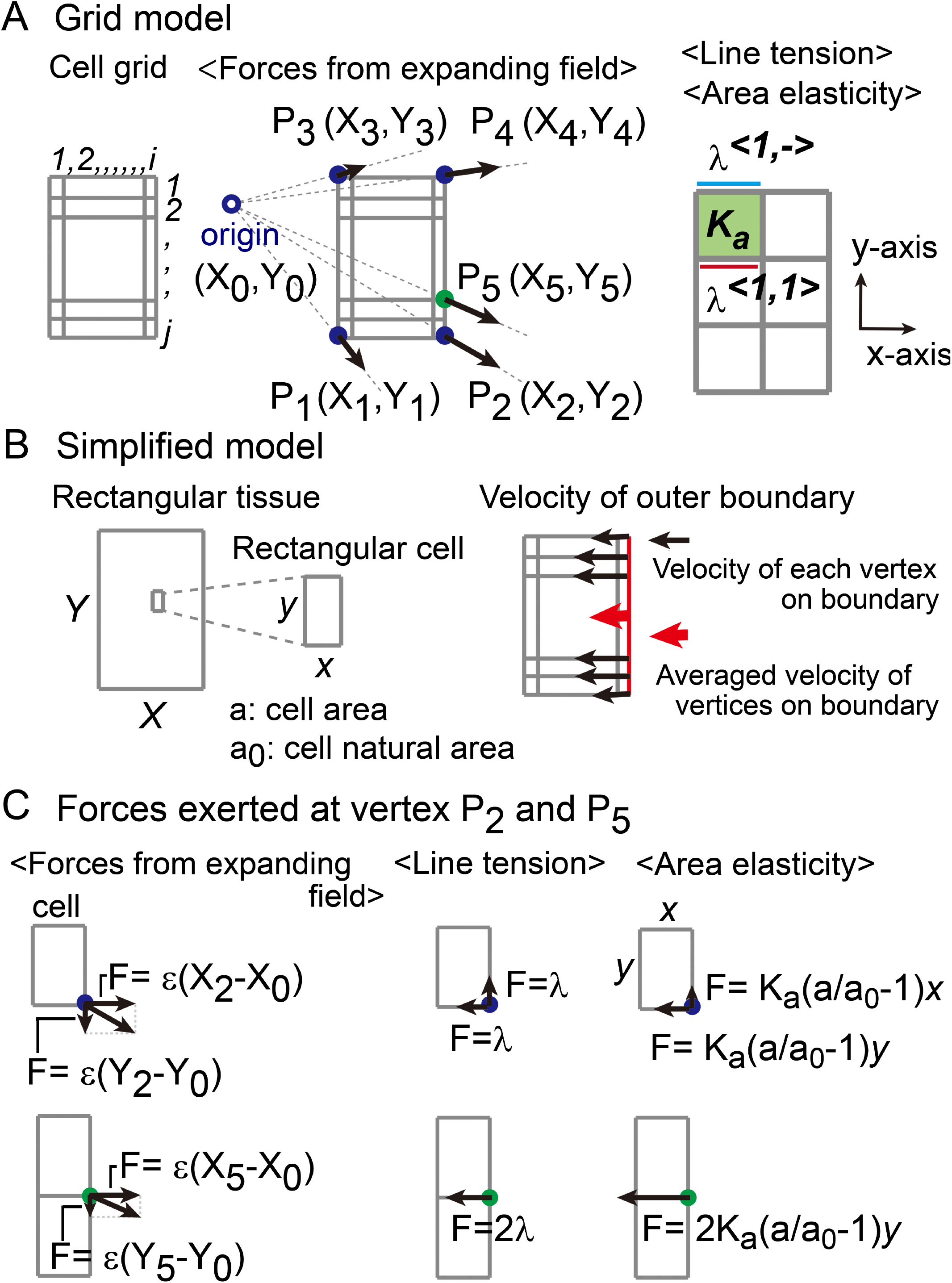
Cell-grid model and its simplification. A) Overview of the cell-grid model. A rectangular cell cluster (P_1_–P_2_–P_4_–P_3_, blue filled circles) that contains rectangular cells (*i×j*) is assumed. Apparent forces provided from field expansion are indicated as arrows at each point of the cell cluster, from P_1_ to P_5_. Line tension *λ* between cells (*λ*^<1,1>^) and on boundaries (*λ*^<1,−>^), along with the area elasticity (*K_a_*), are shown. B) Model simplification. The cell cluster is fixed as rectangular. Edge lengths are *X* and *Y*. Cell shapes are determined as described in Figure S2. The velocity (red arrow) of the right-side outer boundary (red line) is shown. Cells are shown with edge lengths *x* and *y*, the area of cell(*a*), and the preferred area of cell (*a_0_*). The velocity (black arrows) of each vertex was calculated from the values of the forces at each vertex, as shown in C. The velocities of the four lines of the outer boundary are assumed to be the averaged velocity of vertices on each line (Fig. S2). C) Force components exerted on vertices. Filled circles correspond to P_2_ (blue) and P_5_ (green), shown in A. Forces are derived from field expansion, line tension, and area elasticity. The forces exerted at P_2_ or P_5_ are shown along the x- and y-axes.

We then further simplified the grid model by assuming that a cell cluster in the grid was fixed as rectangular, and then considered the dynamics of the outer boundary of the cluster (Fig. 3B and S2). The edge lengths of the cluster were defined as *X* and *Y* along the x- and y-axes, respectively; the dynamics of the inner region were not directly considered. Cell shapes were assumed to be identical. In other words, when *X*, *Y*, *i*, and *j* are given, cell shapes are geometrically determined as follows: cellular edge lengths are *x* = *X/i* and *y* = *Y/j* (Fig. 3B, left panel and S2). Therefore, in our model, when *X* and *Y* temporally evolve during simulations, the cell shapes are automatically determined. The forces exerted on each vertex of the outer boundary were written as shown in Figure 3C (the coordinates of the origin, vertex P_2_, and vertex P_5_ were defined in Figure 3A). The velocities of the four outer boundary lines were defined as the average velocity of the vertices on each line (Fig. 3B, right panel and S2). Consequently, we obtained a simplified model describing the dynamics of the length of the cell cluster along the x- and y-axes, as follows (SI):

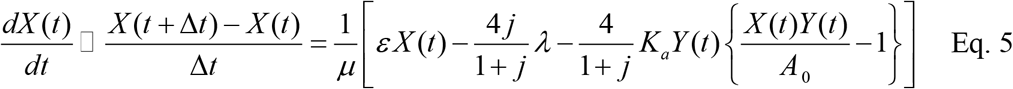

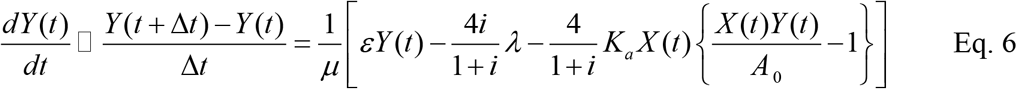

where *A_0_* is the preferred area of the cell cluster (*A_0_* = *i j a_0_*, where *a_0_* is the preferred area of a cell) (Fig. 3B, left panel); *ε* is a coefficient related to field expansion, defined in Equation 4; and *i* and *j* are constants. Importantly, this model contains only two variables, *X* and *Y*. These equations do not contain the coordinates of the origin (*X0, Y0*) or the points on the outer boundaries (*X_1,2,3_,,,, Y_1,2,3_,,,*), indicating analytically that deformation of the cell cluster does not depend on the position of the origin or the cell cluster, consistent with Figures S1C and D in the case of the vertex model.

### Phase plane/space analysis of cell cluster deformation

Prior to our dissection of the roles of the field and cell clusters, we investigated the dynamics of cell cluster shapes (*X* and *Y*) in this simplified model by performing phase plane/space analyses. In the absence of field expansion, a stable equilibrium state was observed (Fig. 4A, right panel, open circle), and most cell clusters were expected to reach this state according to these trajectories (Fig. 4A, right panel, blue and yellow arrows). In the presence of field expansion, tracing of the trajectories led us to expect that most cell clusters would rapidly approach the *dY/dt* or *dX/dt* nullcline, deform along one of the nullclines, and then continued to elongate along the x- or y-axis, respectively (Fig. 4A, left panel, blue and yellow arrows). In addition, we observed no stable equilibrium states in the presence of field expansion, although two saddle states were found. Note that Figure 4A is a representative result under parameter sets in which a stable equilibrium state existed in the absence of field expansion; in biological contexts, stable equilibrium states should exist under no-expansion conditions. Moreover, the field expansion rate was set to be sufficiently strong to induce elongation (large *ε* value). In other parameter sets, various nullclines emerged as shown in Figure S3A, and the combination of various *dX/dt* and *dY/dt* nullclines became very complicated (data not shown). However, such cases are neither important nor realistic in biological contexts, because they do not satisfy the two criteria above: the existence of stable equilibrium states under no-expansion conditions and a high rate of field expansion.

**Figure 4:**
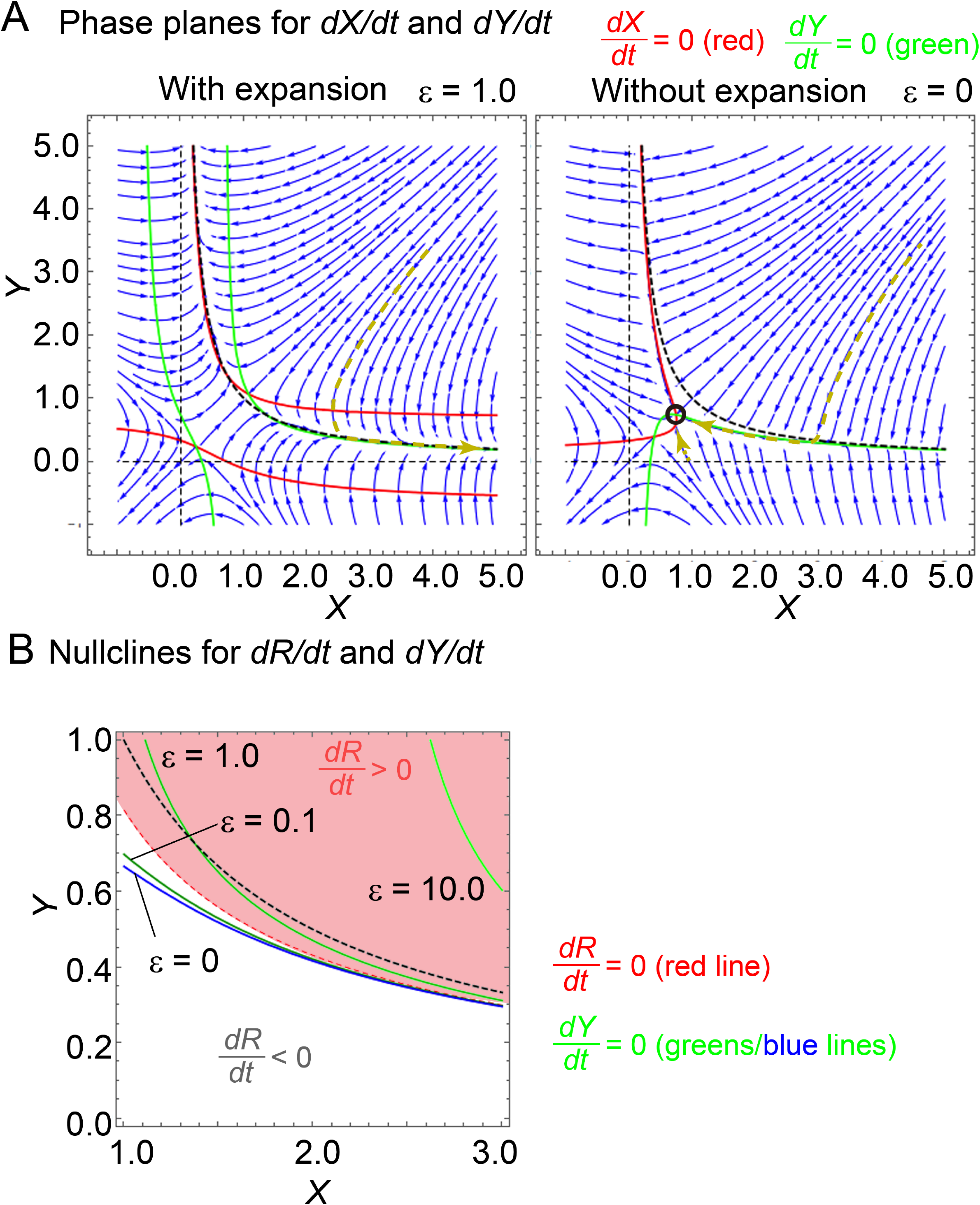
Phase plane analysis of simplified model. A) Phase planes/spaces with (*ε* = 1) or without (*ε* = 0) field expansion. Streams of dynamics and examples of trajectories running along the streams are shown as blue and yellow arrows, respectively. *X* and *Y* are the lengths of the cell cluster along each axis. Red curve, *dX/dt* = 0 (Eq. 5) nullcline; green curve, *dY/dt* = 0 (Eq. 6) nullcline; stable equilibrium state, open circle; black dashed curve, *XY* = *A_0_*. *λ* = 0.2, *K_a_* = 6.0, *a_0_* = 0.1, *i* = 10, *j* = 10, *μ* = 1.0. Plots were generated using the Mathematica software. B) Phase plane with *dR/dt* nullcline. The phase plane in A is enlarged. *dY/dt* = 0 nullclines for each value of *ε* are written next to green or blue curves. *dR/dt* nullclines are shown in red dashed lines. *dR/dt* > 0 regions are colored light red. *XY* = *A_0_* is indicated by a black dashed line.

### Dissection of the roles of cell cluster–intrinsic features and field expansion

To determine whether elongations of cell clusters are caused by the properties of an expanding field or cell cluster, or their combined effects, we directly analyzed the dynamics of the *X-Y* ratio of the cell cluster. By modifying Equations 5 and 6, we obtained a differential equation for the *X-Y* ratio, *R*, as follows (SI);

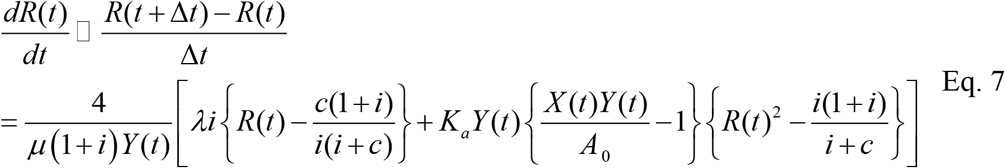

where *c* is the total cell number (*ij*) in the cell cluster. In the specific case where *i* = *j*, the following equation was obtained:

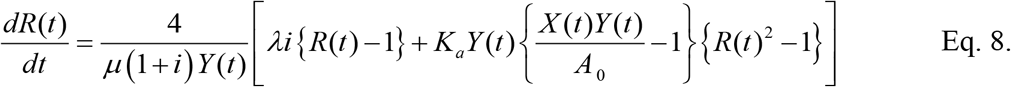

To understand the dynamics of *R*, the sign of this equation is critical. Because all variables [*μ, λ, K_a_, i, X(t), Y(t), R(t)*, and *A_0_*] in Equation 8 are positive, the sign of this equation is largely dependent on the following three terms: {*R*(*t*)-1}, {*R*(*t*)^2^-1}, and {*X(t)Y(t)/A_0_*-1}. For instance, when the cell cluster area [*A(t) = X(t)Y(t)*] is greater than *A_0_*, once *R* becomes larger than 1, it is expected to continuously increase, i.e., the cell cluster is expected to continuously elongate. *A(t)* is expected to become large when field expansion is strong.

Equation 8 includes *λ* and *K_a_*, but, not *ε*, which is related to field expansion. Since *λ* and *K* determines intrinsic features of cells, these facts indicate that the dynamics of *R* are primary dependent on the cell-intrinsic features, but not on field expansion. On the other hand, the field expansion characterized by *ε* may indirectly affect the dynamics of *R*, likely through an increase in *A(t)*, which is critical for the sign of Equation 8 as described in the previous paragraph.

To support these ideas, we plotted the nullcline of *dR/dt* on the *X-Y* phase plane (Fig. 4B and S3B, red dashed lines). The position of the nullcline is absolutely the same regardless of the presence of field expansion, as expected from Equations 7 and 8, indicating that the nullcline position is determined by the cell cluster–intrinsic features, but not by field expansion. Furthermore, when we focused on the case in which *X* > *Y* (the lower region of the line of *Y* = *X*, in which the entire area of Figure 4B is included), the region with *dR/dt* > 0 is located around the region where the cell clusters are enlarged, which corresponds to the upper right region of Figure 4B as well as of the *X-Y* plane, where *X(t)Y(t)* is greater. This indicates that the enlarged cell cluster has an intrinsic feature that increases its own asymmetry. Thus, elongation of the cell clusters is primarily caused by intrinsic features of the enlarged cell clusters, but not the field.

### Theoretical analyses of role of field expansion

Next, we theoretically examined whether the strengths of field expansion applied in our analyses are sufficient to maintain cell clusters under enlarged states with *dR/dt* > 0. The trajectories of cell cluster deformation on the *X-Y* phase plane are expected to approach the equilibrium condition *dX/dt* (= 0) or *dY/dt* (= 0), and to move along these nullclines (Fig. 4A). According to Figure 4A, when we focused on the case of *X* > *Y*, the *dY/dt* nullcline seemed to be responsible for the trajectories. Application of field expansion caused the *dY/dt* nullcline to shift upward along the y-axis (Fig. 4B), corresponding to the increase in *Y*, and thus to enlargement of the cell clusters. We compared the location of the *dY/dt* nullcline with that of the *dR/dt* nullcline. Under stronger field expansion conditions (*ε* ≥ 1), the *dY/dt* nullcline becomes located in the region with *dR/dt* > 0, which should lead to cell cluster elongation (Fig. 4B). This is consistent with the outcomes in the left panel of Figure 4A, where *ε* was set as 1. Taken together, these findings indicate that field expansion considered in our analyses (*ε* ≥ 1) is sufficiently strong for the cell clusters to manifest their intrinsic features that cause them to elongate. Consequently, the effect of field expansion on cell cluster elongation is derived from enlargement of cell clusters.

### Origin of cell cluster–intrinsic feature involved in elongation

What is the source of the cell cluster–intrinsic feature responsible for elongation? To address this question, we considered enlarged cell clusters and focused on the forces derived from the line tensions and area elasticities. To calculate the contributions of these forces to the rates of change in *X* and *Y* in enlarged cell clusters, we extracted the effects of these forces from Equations 5 and 6, which define the rates *dX/dt* and *dY/dt*. If cell numbers *i* and *j* were assumed to be sufficiently greater than 1, the deformation rates along x- and y-axes can be obtained as follows (SI, Method of calculating deformation rates provided from cell cluster–intrinsic forces):

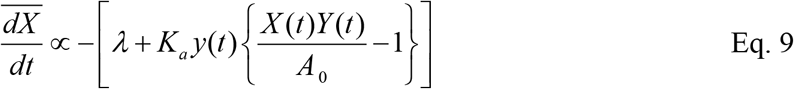

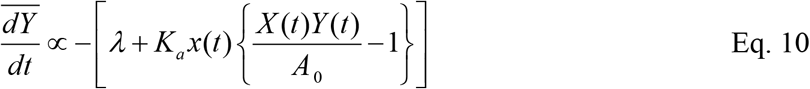

Here, 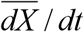 and 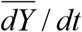 are the deformation rates provided from the forces derived from the line tensions and area elasticities. *x(t)* and *y(t)* are the lengths of cell edges along the x- and y-axes, as previously defined. We considered various shapes of cell clusters, and compared the two rates against each other (Fig. 5 and S4). The two rates are schematically shown as absolute values (*ΔX* and *ΔY*) in Figure 5 and S4.

**Figure 5:**
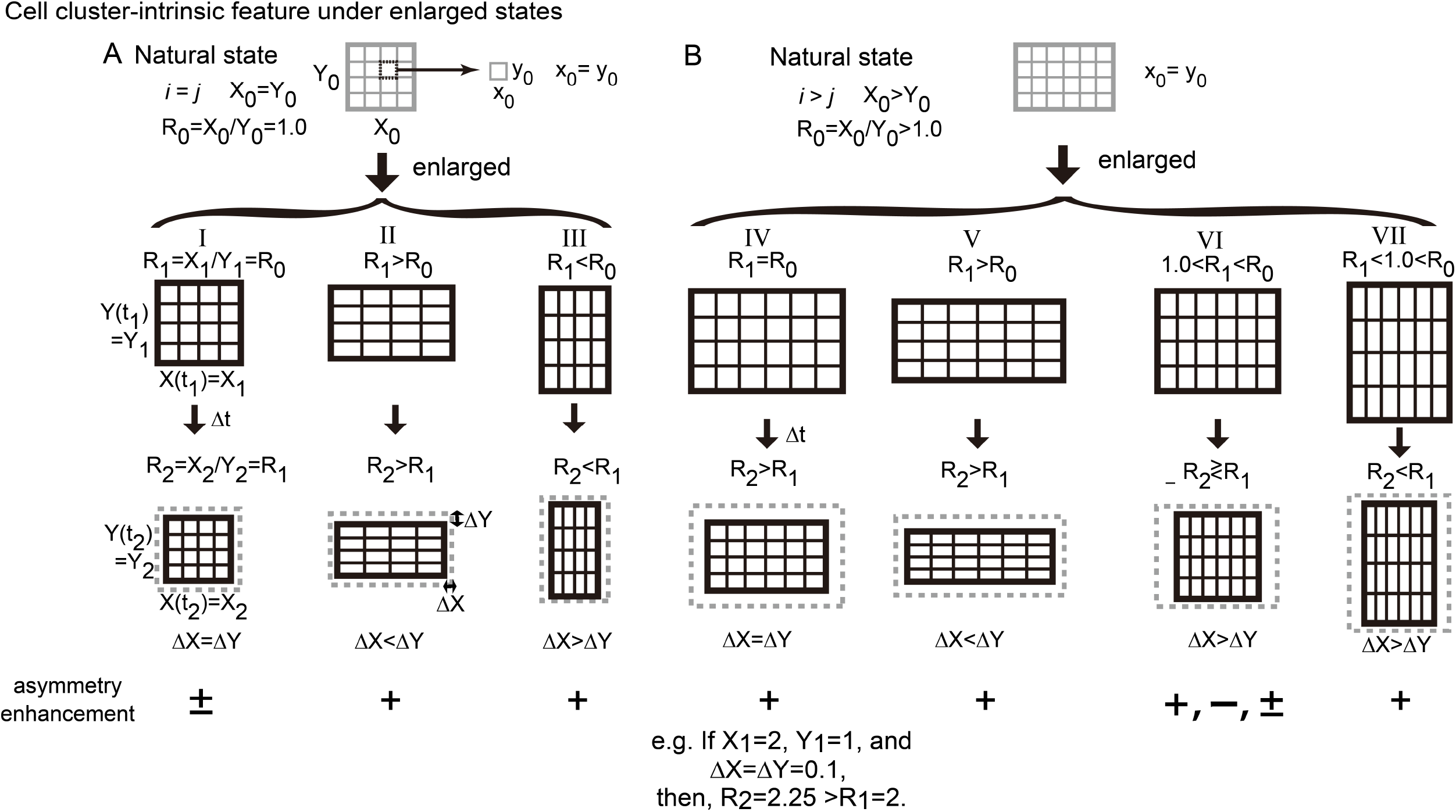
Cell cluster–intrinsic feature that causes elongation. Cell cluster–intrinsic feature is schematically represented for the case of an enlarged state. The natural states of cell clusters are square or rectangular, as indicated by gray grids in the top panels (A and B, respectively). *X_0_* and *Y_0_* are the natural edge lengths of the cell clusters. *i* and *j* are the cell numbers along the x- and y-axes, respectively. Whether various enlarged cell clusters (black grids) enhance their own asymmetries during a short time period Δ*t* is indicated in the bottom panels, in which the shapes prior to shrinkage are described as gray dashed rectangles (I–VII). Time points before and after shrinkage are *t_1_* and *t_2_*, respectively (*t_2_* = *t_1_* + *Δt*). The edge lengths of the cell clusters before deformation are *X_1_* and *Y_1_*, and the lengths after deformation are *X_2_* and *Y_2_*. An example of the dynamics of *R* was calculated. Displacements of the outer boundaries are Δ*X* and Δ*Y*, where Δ*X* = |*X_2_* – *X_1_*|/*2* and Δ*Y* = |*Y_2_* – *Y_1_*|/*2*. The deformations were calculated by considering the line tensions and area elasticities, as shown in Equations 9 and 10. In cases II, IV, and V, the shapes elongated that are along the x-axis will be more elongated, enhancing the asymmetry (asymmetry enhancement is “+”), and *R* will be elevated. In cases III and VII, the shapes that are elongated along the y-axis will be more elongated along the y-axis (asymmetry enhancement is “+”), and *R* will be reduced. In case I, *R* will not be changed (asymmetry enhancement is “±”), but if a slight change in *R* is introduced due to fluctuation of forces, external perturbations, or other factors, the cell cluster will convert to cases II or III, resulting in elongation. In case VI, whether *R* will be increased or decreased depends on the contribution ratio between the area elasticity and the line tension (asymmetry enhancement is “+”, “-”, or “±”). In summary, the cell clusters will increase their asymmetries except in some instances of case VI. In case IV, values of deformation are exemplified. In this condition, *X_2_* = *X_1_* – 2Δ*X* = 1.8, and *Y_2_* = *Y_1_* – 2Δ*Y* = 0.8, and thus, *R_2_* = *X_2_/Y_2_* = 2.25.

In enlarged cell clusters (*X(t)Y(t)*>*A_0_*), the signs of the two deformation rates are negative, indicating shrinkage along the x- and y-axes. When each cell is square (*x(t) = y(t)*), the shrinkage rates along the two axes are the same (Fig. 5-I and IV; *△X=△Y*,). Notwithstanding this situation, enlarged cell clusters with *X* > *Y* are expected to increase *R* (*X-Y* ratio), as exemplified in Figure 5-IV. This feature is provided from both the line tensions (*λ*) and the area elasticities (*K_a_*), as speculated based on Equations 9 and 10. Moreover, in the case that *x(t) > y(t)*, the shrinkage rate is higher along the y-axis, and enlarged cell clusters with *X* > *Y* dramatically increase *R* (Fig. 5-II and V; *ΔX* < *ΔY*). This feature is largely dependent on the area elasticities. Essentially, when the elongation directions of a cell cluster and each cell are correlated with each other, the cell cluster is expected to elongate (Fig. 5-II, III, V, and VII). Under non-correlated conditions, the outcomes vary (Fig. 5-VI). By contrast, shrunken cell clusters (*X(t)Y(t)<A_0_*) have reversed features that reduce their asymmetries (Fig. S5). Thus, although the mean mechanical properties of cells are symmetric under normal conditions [*X(t)Y(t) ≅ A_0_*], cell clusters can induce their asymmetric deformation in enlarged states (Fig. 6). In conclusion, under stronger field expansion, cell clusters are maintained in an enlarged state that has an intrinsic feature that increases the asymmetry of the cluster, resulting in elongation.

**Figure 6:**
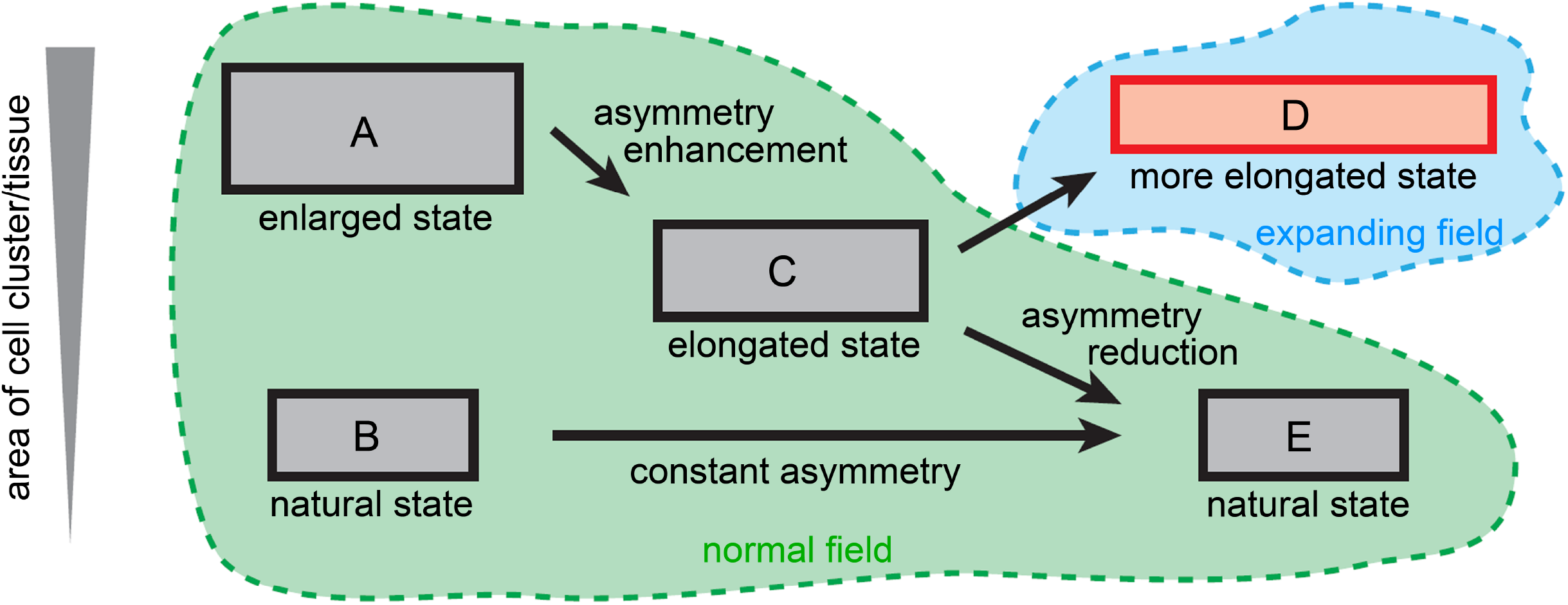
Mechanism of cell cluster elongation. Schematic representation of the dynamics of cell cluster or tissue deformation under an expanding field and a normal field. Cell clusters are depicted by gray and red rectangles. The areas of the cell clusters increase toward the upper regions of the figure, as shown by the indicator at left. The expanding field is depicted in light blue, and the normal field in green. Under a normal field, a cell cluster in the natural state (i.e., with an area nearly equal to its natural area) does not deform and consequently retains its asymmetric shape (B and E). An enlarged cell cluster has intrinsic features that enhance its asymmetric shape, resulting in elongation even under a normal field (A and C). However, when the area becomes nearly equal to the natural area, the effects of enhancement and reduction of asymmetry balance each other (Fig. S4), resulting in convergence to the natural shape (C and E). When a cell cluster is maintained in an enlarged state under an expanding field, it continuously increases its asymmetry, resulting in more and more elongation (D).

## Discussion

In this study, we theoretically investigated the effect of isotropically expanding fields on morphogenesis of tissues and cells located on those fields. Although our study was initially inspired by elongation of the mouse notochord, isotropic expansion of fields/tissues occurs frequently in multiple developmental processes, so we developed a generalized theoretical model. Therefore, our analyses were performed under very simplified and ideal conditions. Conversely, for individual cases such as the mouse notochord, the contributions of field expansion will be quantitatively validated by further comparisons of experimental and theoretical analyses.

Using the generalized model, we found that cell clusters modeled by a standard multi-cellular framework elongated on isotropically expanding fields, although neither the cell clusters nor the field have explicit features providing asymmetries or anisotropies, except for the initial cell configurations. These isotropically expanding fields were also involved in elongation of cells and their directional alignment, consistent with the development of the mouse notochord. We then dissected the contributions of cell clusters and field expansion to elongation through the phase plane analyses and subsequent theoretical analyses, and established a theoretical explanation for elongation of cell clusters. Elongation of cell clusters is primarily caused by a cell cluster–intrinsic feature that enhances their asymmetry (Fig. 6A and C), which is hidden under the no–field expansion condition (Fig. 6B and E). Expansion of a field causes a cell cluster to manifest its intrinsic features by maintaining it in an enlarged state (Fig. 6D). Usually, the direction of cell cluster deformation is correlated with that of extrinsic forces (6, 7). However, on isotropically expanding fields, cell clusters do not deform in response to an extrinsic cue, but along a direction determined by their intrinsic feature, yielding a counterintuitive outcome. Field expansion maintains cells and tissues under a non-equilibrium condition by enlarging them, leading to collective cell patterning and tissue elongation (Fig. 6D). These findings reveal a novel cooperative relationship between cell clusters or tissues and extrinsic cues, and the substantial contributions of expanding fields/environments to cell and tissue morphogenesis.

Elongation of cell clusters such as the notochord were previously understood in the context of cell-autonomous movements accompanying cell intercalations and subsequent convergent extension processes (1). We cannot rule out the possibility that cell-autonomous movements are involved in elongation of the mouse notochord. However, the direction of cell elongation was not consistent with the previous model based on the cell-autonomous movements; moreover, elongated tissue with a single-cell width could only be achieved using field expansion conditions, but not the previous model (data not shown). Accordingly, we favor the idea that the mouse notochord is primary driven by non–cell-autonomous movements, at least during the developmental stages that we observed.

Although it is possible that our results are specific to our models, we did test two models (vertex and grid). Moreover, when a cell was modeled as a particle, and line tensions and area elasticity were not implemented (21), cell clusters also elongated (data not shown). A physical experiment using metal beads placed on an expanding latex membrane also yielded elongation, as previously reported (13). Therefore, we believe that our results are not model-specific. In addition, three-dimensional isotropic expansion occurs in the universe, in which galaxies tend to form elongated/filamentous superclusters (22). These observations imply the generality of symmetry-breaking based on our model in tissues, cells, and physical phenomena.

Expansion of fields is a common process in growing embryos or tissues that are increasing in size. Volume increases in embryonic cavities contribute to the growth of embryos, and these cavities have been speculated to play a role in the morphogenesis of mice and fishes (16, 23). In growing embryos/tissues, some cell populations contribute to the increase in their sizes through radial interactions, etc. (Fig. 1E), and function as an expanding field, whereas other cells in contact with these populations are moved passively. Epiboly is a basic developmental process during which the area of a cell sheet is increased by radial intercalation (16, 17). Epiboly also occurs in developing skin (24). These examples include not only isotropically but also anisotropically expanding fields; however, the volume increase in embryonic cavities and epiboly can provide almost isotropic expansion of fields. Growth of tubular organs in both the circumferential and longitudinal directions can also provide expanding fields. Because any expanding fields are expected to enlarge cell clusters (Fig. 6D), the cell cluster–intrinsic feature that promotes asymmetric deformation can potentially manifest under multiple situations arising growing tissues. Under these conditions, the direction of deformation of cell clusters is influenced by the anisotropy of the fields, as well as the cell cluster–intrinsic feature, leading to complicated dynamics. Simultaneously, these situations can also modulate the direction of cell elongation and cellular alignment, both of which are important for morphogenesis (6, 25). Consistent with this idea, anisotropic extrinsic forces provided by wing boundaries in *Drosophila* are involved in cell elongation and the alignment direction of PCP (6).

Friction between fields and contacting cells is expected to arise during development of multicellular systems, including germ layers, especially during epiboly (7, 8, 14), the epidermis during pregnancy (26), and possibly epithelial cells in tissues contacting mesenchymal or smooth muscle layers (10, 27). Recently, cell–extracellular matrix (ECM) interactions, which would be related to the frictional forces, have been recognized as important for morphogenesis (28, 29). Therefore, to understand morphogenesis involving epithelial and mesenchymal cells and ECMs, the expanding dynamics of these cells and structures are critical parameters.

Local movement of fields is also possible. Local frictional forces from the mesoderm are exerted during epiboly in zebrafish, which is critical for the placement of neural cell populations in the ectoderm (8). Cell clusters sometimes exhibit distorted shapes during development (4, 30, 31) and in the skin of adult chimeric animals, similar to a shape obtained by our simulation (Fig. 5B, top panel). Furthermore, in combination with other components such as local cell–field adhesion, more complicated patterns are formed in fish (14). The effects of fields on cell movements have not been extensively studied, but the dynamics of fields may play important roles in the elongation of cell populations. The results of this study provide insight into the potential of field dynamics.

## Author contributions

H.K. designed and performed the research; H.K. and T.F. wrote the manuscript; and T.F. provided critical suggestions.

## Acknowledgements

We thank Drs. Tatsuo Shibata and Yohei Kondo for critical reading of the manuscript. This work was supported by KAKENHI from the Japan Society for the Promotion of Science (JSPS) for T.F. and H.K., and the National Institutes of Natural Sciences (NINS) program for cross-disciplinary science study for H.K. The authors declare no competing interests.

## References

1. Keller, R., L. Davidson, A. Edlund, T. Elul, M. Ezin, D. Shook, and P. Skoglund. 2000. Mechanisms of convergence and extension by cell intercalation. Phil.Trans. R. Soc. Lond. B. 355: 897–922.

2. Honda, H., T. Nagai, and M. Tanemura. 2008. Two different mechanisms of planar cell intercalation leading to tissue elongation. Dev. Dyn. 237: 1826–1836.

3. Zajac, M., G.L. Jones, and J.A. Glazier. 2000. Model of convergent extension in animal morphogenesis. Phys. Rev. Lett. 85: 2022–2025.

4. Mao, Y., A.L. Tournier, P.A. Bates, J.E. Gale, N. Tapon, and B.J. Thompson. 2011. Planar polarization of the atypical myosin Dachs orients cell divisions in Drosophila. Genes Dev. 25: 131–136.

5. Mc, D. de la L., and T. Bj. 2017. Forces shaping the Drosophila wing. Mech Dev. 144: 23–32.

6. Aigouy, B., R. Farhadifar, D.B. Staple, A. Sagner, J.C. Röper, F. Jülicher, and S. Eaton. 2010. Cell Flow Reorients the Axis of Planar Polarity in the Wing Epithelium of Drosophila. Cell. 142: 773–786.

7. Butler, L.C., G.B. Blanchard, A.J. Kabla, N.J. Lawrence, D.P. Welchman, L. Mahadevan, R.J. Adams, and B. Sanson. 2009. Cell shape changes indicate a role for extrinsic tensile forces in Drosophila germ-band extension. Nat. Cell Biol. 11: 859–864.

8. Smutny, M., Z. Ákos, S. Grigolon, S. Shamipour, V. Ruprecht, D. Čapek, M. Behrndt, E. Papusheva, M. Tada, B. Hof, T. Vicsek, G. Salbreux, and C.P. Heisenberg. 2017. Friction forces position the neural anlage. Nat. Cell Biol. 19: 306–317.

9. Sugimura, K., and S. Ishihara. 2013. The mechanical anisotropy in a tissue promotes ordering in hexagonal cell packing. Development. 140: 4091–4101.

10. Koyama, H., D. Shi, M. Suzuki, N. Ueno, T. Uemura, and T. Fujimori. 2016. Mechanical Regulation of Three-Dimensional Epithelial Fold Pattern Formation in the Mouse Oviduct. Biophys. J. 111: 650–665.

11. Shindo, A., T.S. Yamamoto, and N. Ueno. 2008. Coordination of cell polarity during Xenopus gastrulation. PLoS One. 3: e1600.

12. Wallingford, J.B. 2012. Planar Cell Polarity and the Developmental Control of Cell Behavior in Vertebrate Embryos. Annu. Rev. Cell Dev. Biol. 28: 627–653.

13. Imuta, Y., H. Koyama, D. Shi, M. Eiraku, T. Fujimori, and H. Sasaki. 2014. Mechanical control of notochord morphogenesis by extra-embryonic tissues in mouse embryos. Mech. Dev. 132: 44–58.

14. Reig, G., M. Cerda, N. Sepúlveda, D. Lores, V. Castañeda, M. Tada, S. Härtel, and M.L. Concha. 2017. Extra-embryonic tissue spreading directs early embryo morphogenesis in killifish. Nat. Commun. 8: 15431.

15. Morishita, Y., and T. Suzuki. 2014. Bayesian inference of whole-organ deformation dynamics from limited space-time point data. J. Theor. Biol. 357: 74–85.

16. Trinkaus, J.P. 1984. Cells into Organs. The Forces That Shape the Embryo. Prentice-Hall Inc.

17. Solnica-Krezel, L. 2006. Gastrulation in zebrafish — all just about adhesion ? Curr. Opin. Genet. Dev. 16: 433–441.

18. Fletcher, A.G., M. Osterfield, R.E. Baker, and S.Y. Shvartsman. 2014. Vertex models of epithelial morphogenesis. Biophys. J. 106: 2291–2304.

19. Suzuki, M., M. Sato, H. Koyama, Y. Hara, K. Hayashi, N. Yasue, H. Imamura, T. Fujimori, T. Nagai, R.E. Campbell, and N. Ueno. 2017. Distinct intracellular Ca 2 + dynamics regulate apical constriction and differentially contribute to neural tube closure. Development. 144: 1307–1316.

20. Okuda, S., Y. Inoue, M. Eiraku, T. Adachi, and Y. Sasai. 2014. Vertex dynamics simulations of viscosity-dependent deformation during tissue morphogenesis. Biomech. Model. Mechanobiol. 14: 413–425.

21. Merkel, M., and M.L. Manning. 2017. Using cell deformation and motion to predict forces and collective behavior in morphogenesis. Semin. Cell Dev. Biol. 67: 161–169.

22. Tully, R.B., H. Courtois, Y. Hoffman, and D. Pomarède. 2014. The Laniakea supercluster of galaxies. Nature. 513: 71–73.

23. Tam, P.P.L., and R.R. Behringer. 1997. Mouse gastrulation: The formation of a mammalian body plan. Mech. Dev. 68: 3–25.

24. Panousopoulou, E., C. Hobbs, I. Mason, J.B.A. Green, and C.J. Formstone. 2016. Epiboly generates the epidermal basal monolayer and spreads the nascent mammalian skin to enclose the embryonic body. J. Cell Sci. 129: 1915–1927.

25. Koyama, H., and T. Fujimori. 2018. Biomechanics of epithelial fold pattern formation in the mouse female reproductive tract. Curr. Opin. Genet. Dev. 51: 59–66.

26. Ichijo, R., H. Kobayashi, S. Yoneda, Y. Iizuka, H. Kubo, S. Matsumura, S. Kitano, H. Miyachi, T. Honda, and F. Toyoshima. 2017. Tbx3-dependent amplifying stem cell progeny drives interfollicular epidermal expansion during pregnancy and regeneration. Nat. Commun. 8: 508.

27. Shyer, A.E., T. Tallinen, N.L. Nerurkar, Z. Wei, E.S. Gil, D.L. Kaplan, C.J. Tabin, and L. Mahadevan. 2013. Villification: how the gut gets its villi. Science. 342: 212–8.

28. Goodwin, K., S.J. Ellis, E. Lostchuck, T. Zulueta-coarasa, R. Fernandez-gonzalez, K. Goodwin, S.J. Ellis, E. Lostchuck, and T. Zulueta-coarasa. 2016. Basal Cell-Extracellular Matrix Adhesion Regulates Force Transmission during Tissue Morphogenesis. Dev. Cell. 39: 611–625.

29. Ryan, P.L., R.A. Foty, J. Kohn, and M.S. Steinberg. 2001. Tissue spreading on implantable substrates is a competitive outcome of cell-cell vs. cell-substratum adhesivity. Proc. Natl. Acad. Sci. U. S. A. 98: 4323–4327.

30. Vogt, W. 1929. Gestaltungsanalyse am Amphibienkeim mit ortlicher Vitalfarbung. II. Teil: Gastrulation und Mesodermbildung bei Urodelen und Anuren. Roux’ Arch. 120: 384–706.

31. Shi, D., K. Komatsu, M. Hirao, Y. Toyooka, H. Koyama, F. Tissir, A.M. Goffinet, T. Uemura, and T. Fujimori. 2014. Celsr1 is required for the generation of polarity at multiple levels of the mouse oviduct. Development. 141: 4558–4568.

